# Dimension Reduction by Spatial Components Analysis Improves Pattern Detection in Multivariate Spatial Data

**DOI:** 10.1101/2023.10.12.562016

**Authors:** Niklas Kleinenkuhnen, David Köhler, Till Baar, Chrysa Nikopoulou, Vangelis Kondylis, Matthias Schmid, Peter Tessarz, Achim Tresch

## Abstract

We introduce a multivariate statistical approach for pattern recognition in spatial transcriptomics data. Our algorithm (SPACO) constructs a low-dimensional projection of the data maximising Moran’s I, which mitigates non-spatial variation and outperforms PCA for pre-processing. Our method also provides a calibrated, powerful test of spatial gene expression that excels in robustness and specificity.

## Main

Technological advances have made spatially resolved transcriptomics widely applicable in the life sciences, providing near-cellular resolution of tissue heterogeneity and organisation. Neighbouring cells in a tissue are typically of similar types, leading to a dependence of observations on proximity. This has spurred research on statistical strategies for handling the rich data from spatial omics, in particular for the detection of spatially variable genes (SVG) and the digital dissection of tissue into functional regions defined by SVG expression.

Various statistics have been used for SVG detection, including Moran’s I, Geary’s C, Gaussian process models, and non-parametric models. Existing SVG identification methods often test spatial dependence on a per-gene basis^1^. However, gene expression is typically organised in co-regulated networks. To address this, methods like PCA are often used for noise reduction and feature extraction, finding covariance structure in the data, and maximising the variance of the projected data for the first few principal components.

We introduce Spatial Component Analysis (SPACO, Online Methods), a proximity-aware kernel method for spatial data. By replacing PCA’s global variance target with Moran’s I, a measure of local (co)variance, SPACO constructs an ordered sequence of basis vectors, the spatial components (SpaC). Orthogonal data projection onto the first *k* SpaCs maximises Moran’s I, thereby pooling evidence of spatial dependence across genes with similar patterns. This enhances the sensitivity and spatial precision of the signal (Figure 1A).

**Figure 1.**
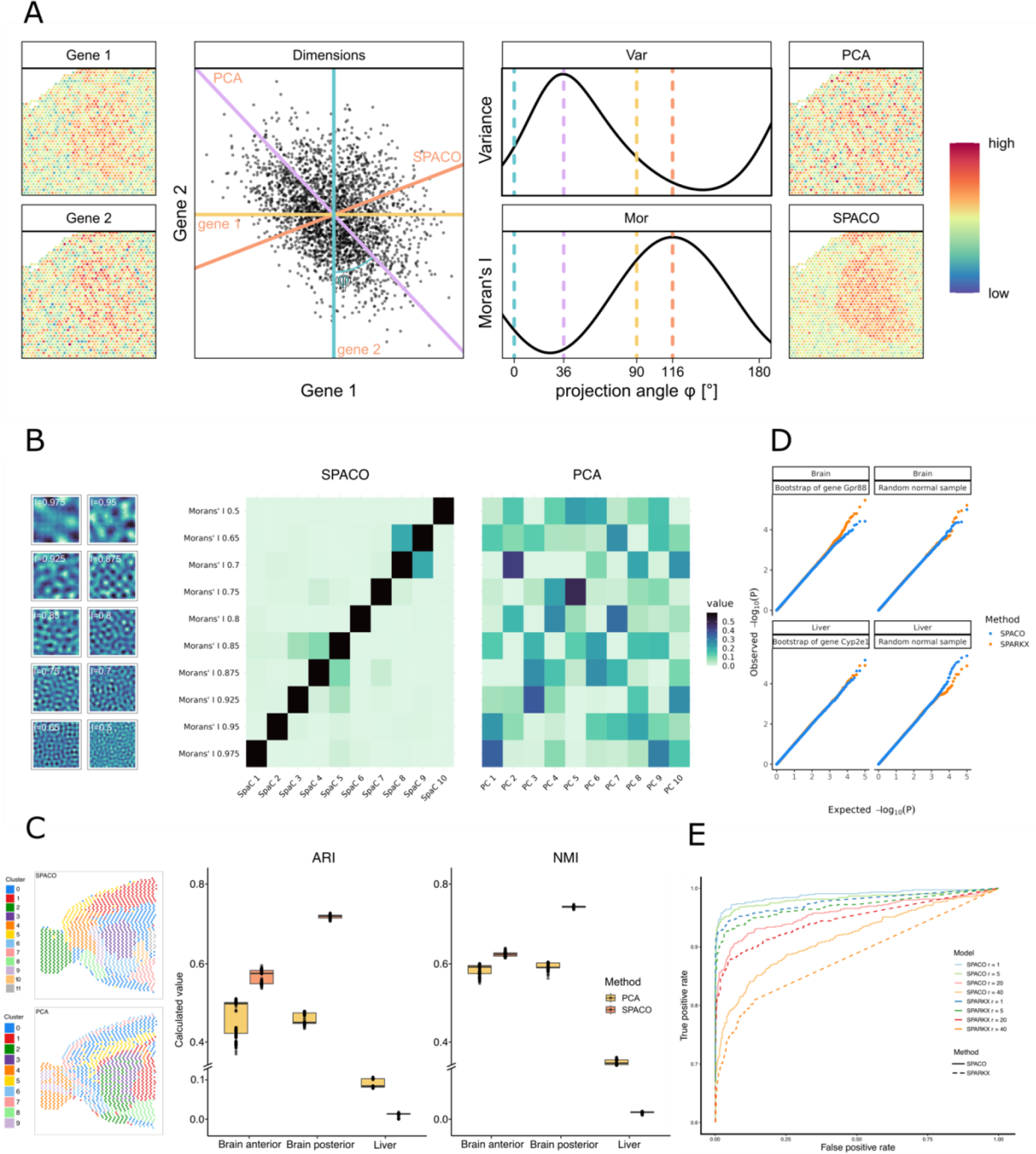
A) Schematic of SPACO and PCA for two genes with weakly spatial patterns (left). Each spot carries two normalised expression values for Gene 1 and Gene 2, shown in the scatterplot. Coloured lines correspond to 1-dimensional subspaces onto which the data is projected. For each projection angle, the variance and Moran’s I of the projected data is shown (top resp. bottom curve). Coloured dotted lines correspond to the projections in the scatterplot. PCA chooses the projection to maximum variance as the first PC (top right), while SPACO chooses the projection with maximum Moran’s I as the first SpaC (bottom right). B) Application of SPACO respectively PCA to ten synthetic patterns with decreasing spatial structure (left, I = Moran’s I). Heatmaps show the squared loadings of each pattern with respect to each base vector (left: SPACO, right: PCA). C) Clusterings following dimension reduction with SPACO (top left), respectively PCA (bottom left). Cluster similarity as measured by adjusted Rand index (Ari, middle) and normalised mutual information (NMI, right) for three representative datasets (brain anterior, brain posterior, liver) using SPACO (red boxes) or PCA (orange boxes) for 1000 initialisations of Louvain clustering. D) Calibration plots of observed (y-axis) versus expected (x-axis) p-values from SPACO (blue) and SPARKX (orange) in the brain (top) and liver (bottom) data in two simulation settings using the null model (left) and random spot permutations of two genes with strong spatial patterns (Gpr88, Cyp2e1). E) ROC curves for SPACO (solid lines) and SPARK (dotted lines) using the anterior brain data and 757 published spatial genes^3^ as cases and 757 randomly permuted copies as controls. Line colours correspond to the increasing difficulty of the task, i.e., noise levels added to the cases.

We demonstrated SPACO’s superiority over PCA as a dimension reduction method by running the Louvain clustering protocol in Seurat^2^, with PCA replaced by SPACO. The clusters resulting from SPACO were visually more coherent (Figure 1C). This was systematically evaluated using two mouse brain and one liver datasets (Online Methods). The spot grid was split into two mutually adjacent sets, shifted by one unit (Supplementary Figure S1A). Each split dataset underwent entirely independent processing, and the adjusted Rand index and normalised mutual information were used to assess the consistency of the two resulting clusterings. SPACO displayed better reproducibility in the brain data (Figure 1C). The performance of SPACO and PCA decreased significantly with the liver data, where the spatial signal is very fine-grained and was essentially eliminated when the data was split.

We compared SPACO and PCA using synthetic genes with varying spatial dependence (Figure 1B, Online Methods). Unlike PCA, the SpaCs aligned well with these genes, demonstrating its expedience as a spatial-aware dimension reduction method. We, therefore, project patterns onto the first few relevant SpaCs. Their number can be determined by a bootstrap procedure (Supplementary Figure S1E, Online Methods) and, unlike PCA, does not need to be determined manually.

We use the length of the projected pattern of a gene as a test statistic for spatial dependence (Online Methods), resulting in a weighted sum of chi-squares as null statistic. We verified the p-value calibration of SPACO and a reference method, SPARKX^4^, under our null model and evaluated their robustness against normality assumption violations (Figure 1D).

We developed a coverage-adjusted local resampling procedure (Online Methods, Figure S1C and D) to gradually reduce the spatial signal while preserving a gene’s relative abundance distribution. Through ROC analysis, we compared SPACO’s specificity and sensitivity to SPARKX, using 757 published spatial marker genes from the mouse brain sagittal-anterior dataset^3^ and their non-spatial twins obtained via random resampling. The difficulty of the SVG detection task was increased stepwise by perturbing the marker genes by local resampling (Fig. 1E). SPACO consistently outperformed SPARKX across all scenarios. Notably, SPARKX found about 15,000 significant SVGs compared to SPACO’s 1,500–2,500. This discrepancy is explained by the sensitivity of SPARKX to a general anticorrelation of non-spatial genes and the spatial pattern of spot-wise coverage (Online Methods, Supplementary Figure S1B).

We show further that for significant SVGs by SPACO, their projection onto relevant SpaCs is actually a denoising operation. The projections resemble their unperturbed original more closely than the measurements (Figure 2A).

**Figure 2.**
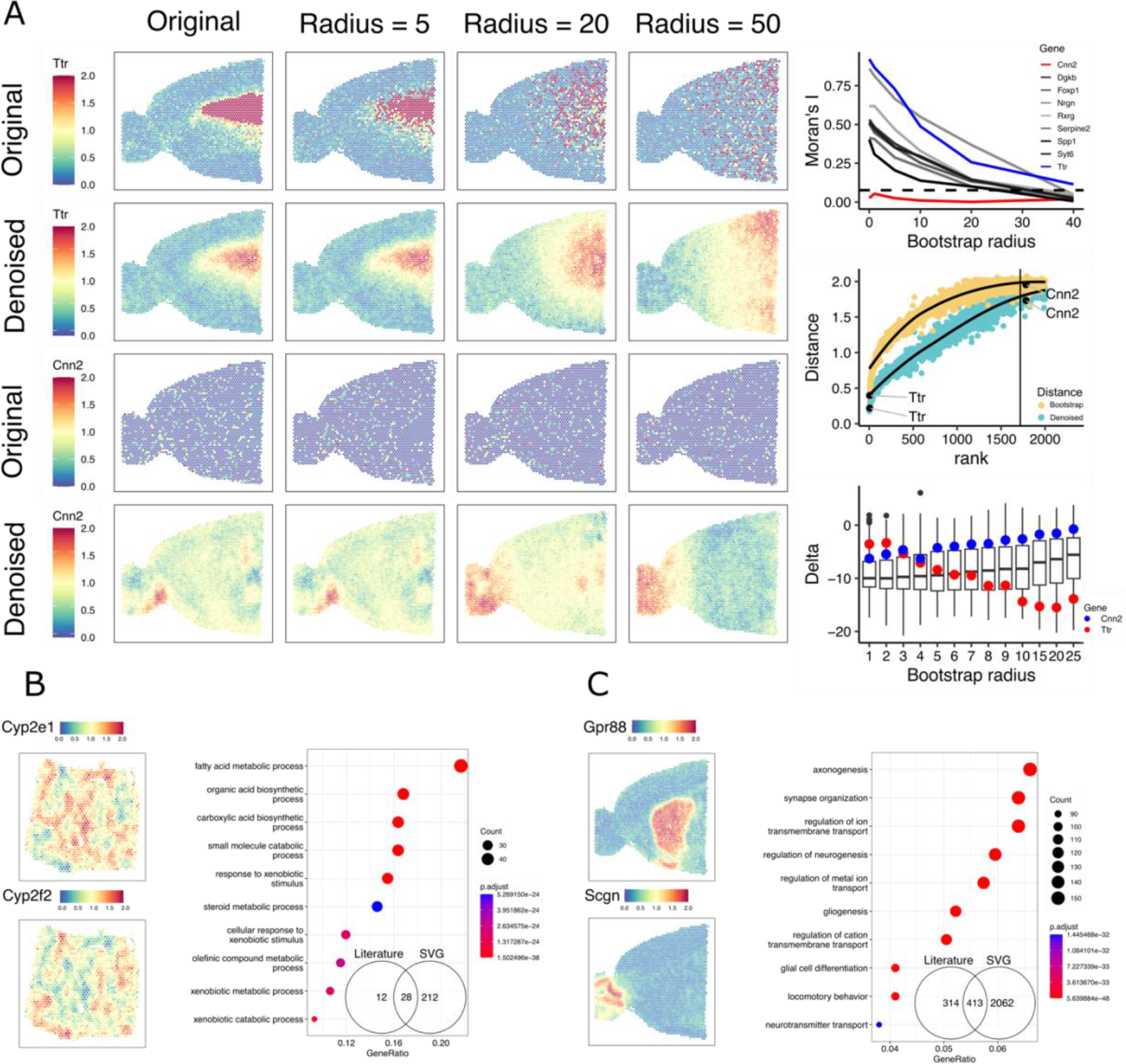
A) Denoising of expression patterns by SPACO. Left: Shown is a gene with a strong spatial pattern (Ttr, 1^st^ row), a gene without a pattern (Cnn2, 3^rd^ row) and their respective SPACO projections (2^nd^ and 4^th^ row). The columns correspond to different radii of local resampling applied to the original measurements. Top right: Decrease of Moran’s I with increasing levels of noise (radius 0 = original data, radius 40 = 75% slide size) for a collection of genes with strong spatial patterns, Ttr (blue) and Cnn2 (red) in particular. The dotted line is the median Moran’s I of all randomly permuted patterns. Middle: Distance (y-axis, average residual sum of squares per spot, RSS) of the original data to the perturbed patterns (radius = 5, yellow), respectively, to their denoised counterparts (turquoise), for the first 2000 genes ranked by significance. The vertical line corresponds to an adjusted p-value of 0.05. (RSS, y-axis). Bottom: Distance reduction (y-axis, Delta = RSS denoised – RSS perturbed) achieved by denoising the set of significant SVG (boxes) for each radius (x-axis). The Delta values for Ttr (blue) and Cnn2 (red) are highlighted individually. B) Denoised profiles of well-known spatial genes in the liver and GO term analysis of significant SVG. Inset: Venn diagram of published vs found SVG. C) same as B), for spatial genes Gpr88 and Scgn in the anterior mouse brain.

Using SPACO on liver data, we recover genes with known periportal-pericentral hepatic lobule gradients^5^, identifying 70% of previously detected genes. Undetected SVG (12 in total) were not found merely due to initial low coverage filtering by the *SCTransform* function of Seurat^6^. GO term analysis links SVGs to fatty acid metabolism and catabolic processes, key functions in periportal and pericentral zones (Figure 2 B and C)^5^. A similar 56% SVG recovery was achieved for anterior brain data^3^. GO term analysis of the genes that were called significant but were not listed in the literature were associated with terms of spatialised processes of the tissues analysed (Supplemental Figure S1F).

Our results confirm that SPACO should replace PCA as the dimension reduction method for spatial omics data. SPACO, with its multivariate approach, increases the sensitivity of SVG detection while controlling false positive rates. It is competitive in terms of time and memory consumption (Supplementary Table 5) and provides denoised spatial patterns that enhance downstream tasks such as clustering and tissue dissection.

## Code Availability

An open-source implementation of SPACO is available on GitHub: https://github.com/IMSBCompBio/SpaCo

The repository includes tutorials and example vignettes for reproducing the presented analyses.

## Contributions

A.T. and P.T. conceived the project. A.T., D.K., M.S., N.K. developed and implemented the method. V.K., C.N, and P.T. generated the data, N.K. and D.K. performed the simulations and analysed the data. T.B., N.K. and D.K. generated the figures for the manuscript. A.T. and N.K. wrote the manuscript with input from all authors. Funding Acquisition: A.T., M.S., P.T.

## Acknoledgements

The authors thank Jason Mueller and Mohammad Hussainy for valuable discussions.

