## Supplementary material for "Dimension Reduction by Spatial Components Analysis Improves Pattern Detection in Multivariate Spatial Data": Online Methods

**by**

Niklas Kleinenkuhnen, David Köhler, Till Baar, Chrysa Nikopoulou, Vangelis Kondylis, Matthias Schmid, Peter Tessarz, Achim Tresch

### Spatial Component Analysis (SPACO)

Suppose we have observed a set of $p$ genes for a set of $n$ spots. The data is represented as a spots × genes matrix $X = \left(x_s^g\right) \in \mathbb{R}^{n\times p}$, where $x_s^g$ denotes the expression of gene $g$ in spot $s$. For a gene $g$, we call the column vector $x^g = \left(x_s^g\right)_{s=1,\dots,n}$ its expression pattern. Generally, we call any vector $x \in \mathbb{R}^n$ an expression pattern. Two expression patterns that are identical up to linear transformation are considered biologically indistinguishable. We therefore standardise the gene expression patterns $x^g$, i.e., we centre them to zero mean zero and to unit variance. We use Moran's I to assess the spatial dependence of a pattern $x$, defined as [7]

$$M(x) = \frac{\frac{1}{|W|}\sum_{i,j=1}^{n} w_{ij}\,(x_i - \bar{x})\left(x_j - \bar{x}\right)}{\frac{1}{n}\sum_i (x_i - \bar{x})^2} \tag{1}$$

Here, $\bar{x}$ is the mean of the values in $x$, $0 \neq W = \left(w_{ij}\right) \in \mathbb{R}^{n\times n}$ is a non-negative, symmetric weights matrix with zero diagonal, and $|W| = \sum_{i,j} w_{i,j} > 0$. The weights matrix is often derived from a translation and rotation-invariant kernel applied to the physical distances of the spots, such as a Gaussian kernel with a width that is about the nearest neighbour distance on the spot grid. In our application, we chose $w_{ij}$ as the function indicating if $i$ and $j$ are nearest or two-step neighbours on the spot grid ($w_{ij} = 1$) or not ($w_{ij} = 0$).

Moran's I can assume values between -1 and 1, with large values above 0 indicating spatially correlated expression, values of 0 indicating no spatial correlation, and small values below 0 indicating spatial anticorrelation. The latter will never happen for spatial transcriptomics data, as the whole spot grid would have to align with a local, biological pattern of mutual anticorrelation. For a centred and scaled pattern $x$, we have

$$1 + M(x) = \frac{1}{n}\sum_{i,j=1}^{n} x_i^2 + \frac{1}{|W|}\sum_{i,j=1}^{n} w_{ij}\, x_i x_j \tag{2}$$

therefore, $1 + M(x)$ is a non-negative quadratic form and can be represented as

$$1 + M(x) = x^T L x \tag{3}$$

with $L = \frac{1}{n}E + \frac{1}{|W|}W$ a symmetric, positive semidefinite matrix. The matrix $L$ gives rise to a scalar product and a metric in pattern space,

$$\begin{aligned} \langle x, y\rangle_L &:= x^T L y \quad, x, y \in \mathbb{R}^n \\ \|x\|_L &:= \sqrt{\langle x, x\rangle_L} \quad, x \in \mathbb{R}^n \end{aligned} \tag{4}$$

The key observation is that this scalar product induces a meaningful similarity measure in pattern space. This is most easily seen by realising that for centred patterns $x$,

$$\|x\|_L^2 = \langle x, x\rangle_L = 1 + M(x) \tag{5}$$

I.e., patterns with strong spatial patterns, respectively large Moran's I have greater length, while non-spatial patterns have smaller length.

Having this notion in mind, we perform principal components analysis of the data $X = (x^g, g = 1, \dots, p)$ in the space $(\mathbb{R}^n, \langle .,. \rangle_L)$. In other words, we perform kernel PCA with kernel $L$. This can be done efficiently by performing the eigenvalue decomposition of the matrix $X^T L X$. Let $u_1, u_2, \dots \in \mathbb{R}^p$ the eigenvectors of $X^T L X$ which we call spatial component (SpaC). Let $\mu_1 \geq \mu_2 \geq \dots$ the corresponding eigenvalues and let $v_j = X u_j \in \mathbb{R}^n$ the pattern conjugate to $u_j$. For each $k$, the space $V_k = \langle v_1, \dots, v_k \rangle$ is optimal in preserving spatial information in the following sense:

Let $\rho(V)$ the "worst" (as measured by Moran's I) spatial pattern contained in the space $V \leq \mathbb{R}^n$,

$$\rho(V) \quad = \quad \min_{x\in V\setminus\{0\}} M(x)$$

( 6 )

Then,

$$\rho(V_k) \quad = \quad \max_{V\leq\langle X\rangle,\, \dim V=k} \rho(V)$$

( 7 )

In other words, the spatial pattern of the worst one-dimensional subspace of $V_k$ is best possible. For the proof of Equation 7, We first prove that the space generated by the first $k$ SpaCs, $U_k = \langle u_1, \ldots, u_k\rangle$, satisfies

$$\tilde{\rho}(U_k) \quad = \quad \max_{U\leq\mathbb{R}^p,\, dimU=k} \tilde{\rho}(U)$$

( 8 )

where

$$\tilde{\rho}(U) = \min_{u\in U, \|u\|_2=1} \|Xu\|_L^2$$

( 9 )

It is elementary to verify that $\tilde{\rho}(U_k) = \mu_k,\ k = 1, \ldots, p$. Let $U \leq \mathbb{R}^p$, dim $U = k$. If $u_p \in U$, then $\tilde{\rho}(U) \leq \mu_p \leq \mu_k$. So, assume without loss that $U \cap \langle u_p\rangle = 0$. Let $W = \langle u_p\rangle^{\perp}$ (in the Euclidean metric $\|.\|_2$), and $U' = \left(U + \langle u_p\rangle\right) \cap W$ the orthogonal projection of $U$ onto $W$. Then dim $U' = k$. As $W$ is $X$-invariant, $\dim W = p - 1$, we have $\tilde{\rho}(U') \leq \mu_k$ by induction on $p$ (the case $p = 1$ being trivial). Any $u \in U$ has a representation $u = \alpha u'(u) + \beta u_p$, with $u'(u) \in U'$, $\|u'(u)\|_2 = 1$, and $\alpha, \beta \in \mathbb{R}$. Assuming further $\|u\|_2 = 1$ yields $\alpha^2 + \beta^2 = \|u\|_2^2 = 1$. It follows that

$$\|Xu\|_L^2 = \alpha^2 \left\|Xu'^{(u)}\right\|_L^2 + \beta^2\mu_p \leq (\alpha^2 + \beta^2)\left\|Xu'^{(u)}\right\|_L^2 = \left\|Xu'^{(u)}\right\|_L^2$$

( 10 )

The inequality in Equation 10 holds because $\mu_p$ is the smallest squared length of a vector $\|Xu'\|_L^2$, $\|u'\|_2 = 1$, that can occur. As dim $U =$ dim $U'$, to every $v' \in U', \|v'\|_2 = 1$, there exists a corresponding $u \in U, \|u\|_2 = 1$ such that in the above representation of $u$, $u'(u) = v'$. It follows that

$$\tilde{\rho}(U) = \min_{u\in U, \|u\|_2=1} \|Xu\|_L^2 \overset{(Eq.10)}{\leq} \min_{u\in U, \|u\|_2=1} \left\|Xu'^{(u)}\right\|_L^2 = \min_{v'\in U', \|v'\|_2=1} \|Xv'\|_L^2 = \tilde{\rho}(U') \leq \mu_k$$

( 11 )

which proves Equation 8.

Now consider some $V \leq \langle X \rangle$, $\dim V = k$. Then, $V = \{Xu; u \in U\}$ for a suitable $k$-dimensional space $U \leq \mathbb{R}^p$. This implies that

$$\begin{aligned}
\tilde{\rho}(U) &= \min_{u \in U, \|u\|_2 = 1} \|Xu\|_L^2 \\
&\overset{(Eq.5)}{=} \min_{u \in U, \|u\|_2 = 1} \big(1 + M(Xu)\big) \\
&\overset{(*)}{=} 1 + \min_{u \in U \setminus \{0\}} M(Xu) \\
&= 1 + \min_{x \in V \setminus \{0\}} M(x) \\
&= 1 + \rho(V)
\end{aligned} \tag{12}$$

The equality in (*) holds because of the invariance of Moran's I with respect to (non-zero) scalar multiplication. It follows that

$$\rho(V_k) \overset{(Eq.12)}{=} \tilde{\rho}(U_k) - 1) \overset{(Eq.8)}{\leq} \tilde{\rho}(U) - 1) \overset{(Eq.12)}{=} \rho(V) \tag{13}$$

which concludes the proof of Equation 7.

### Selection of relevant SpaCs and denoising

We choose the largest $k$ such that the eigenvalue of the $k$-th eigenvector is larger than the 95% quantile of the distribution of the largest eigenvalue occurring in a random data set $X'$ in which the spots of $X$ were permuted randomly. The 95% quantile is estimated from 1,000 such permutations. The projection $P = P_k$ is called the SpaCo projection, and its image is called SpaCo space (Supplementary Figure S1E).

### Detection of spatially variable genes (SVG)

The projection $P$ decomposes a pattern $x$ into the projection into SpaCo space, $Px$, and its $L$-orthogonal complement, $(id - P)x$. For a gene that has a non-random spatial pattern (spatially variable gene, SVG), we expect that $x \approx Px$, and therefore $Px$ should be larger than to be expected by chance. To make this notion quantitative, our null model assumes that an expression pattern $Y \in \mathbb{R}^n$ is spatially independent noise,

$$Y' \sim \mathcal{N}(0, I)$$

( 14 )

Let $s_i = \frac{1}{\sqrt{\mu_i}} X v_i \in \mathbb{R}^n$, $i = 1, \ldots, n$, the $L$-orthonormalized expression patterns conjugate to the SpaCs $v_i \in \mathbb{R}^p$ (note that $< s_i, s_j >_L = \frac{1}{\sqrt{\mu_i \mu_j}} v_i^T X^T L X v_j = \frac{1}{\sqrt{\mu_i \mu_j}} \delta_{ij} \mu_j = \delta_{ij}$ by construction of the SpaCs $v_i, v_j$). Then

$$Y = \sum_{i=1}^{n} < s_i, Y >_L \, s_i = \sum_{i=1}^{n} (s_i^T L Y) \, s_i$$

( 15 )

Let $S_k = (s_1, \ldots, s_k) \in \mathbb{R}^{n \times k}$. The squared length of the $L$-orthogonal projection $PY$ of $Y$ onto the first $k$ SpaC patterns is

$$\|PY\|_L^2 \quad = \left\| \sum_{i=1}^{k} (s_i^T L Y) \, s_i \right\|_L^2 = \quad \sum_{i=1}^{k} (s_i^T L Y)^2 = Y^T L^T S_k S_k^T L Y = Y^T \Sigma Y$$

( 16 )

With $\Sigma = L^T S_k S_k^T L$. By the spectral theorem, we decompose $\Sigma$ into

$$\Sigma = Q C Q^T \in \mathbb{R}^{n \times n}$$

( 17 )

with unitary matrix $Q$ and diagonal matrix $C = \text{diag}(c_1, c_2, \ldots., c_n)$. By rotational invariance of the standard multivariate normal distribution, $Z = Q^T Y \sim \mathcal{N}(0, I)$. It follows that $\|PY\|_L^2$ is a weighted sum of independent $\chi^2(1)$ random variables,

$$T := \|PY\|_L^2 \quad = \quad Y^T \Sigma Y = (Q^T Y)^T \cdot C \cdot (Q^T Y) = Z^T C Z = \sum_{j=1}^{n} c_i \, z_i^2 \sim \sum_{i=1}^{n} c_i \, \chi_i^2$$

( 18 )

with $\chi_i^2 \underset{iid}{\sim} \chi^2(1)$, $i = 1, \ldots, n$. Computations can be accelerated slightly as the non-zero coefficients $c_i$ (of which there are at most $k$) can also be obtained as the eigenvalues of $\tilde{\Sigma} =$

$\left(S_k^T L\right)(L^T S_k) = S_k^T \mathrm{diag}\left(\mu_1^2, \ldots, \mu_k^2\right) S_k \in \mathbb{R}^{k \times k}$. Note that the column vectors in $S_k = (s_1, \ldots, s_k)$ are not orthogonal in Euclidean space, and therefore $\tilde{\Sigma}$ is in general not a diagonal matrix.

Given a gene $g$ with corresponding expression pattern $x_g$ (centered to mean 0 and variance 1), our test statistic is

$$t_g := \left\|P x_g\right\|_L^2 = x_g^T \Sigma x_g \tag{19}$$

and we test the one-sided hypothesis

$$p_g := P\left(T \geq t_g\right) \tag{20}$$

### Synthetic data generation

Let $\frac{1}{|W|} W = V^T \Lambda V$ the eigenvalue decomposition the symmetric matrix which is used to define Moran's I for a given neighbourhood weights matrix $W$, with $V$ a unitary matrix $V = (w_1 \ldots w_n)$, $w_j \in \mathbb{R}^n$, and diagonal matrix $\Lambda = \mathrm{diag}(\lambda_1, \ldots, \lambda_n)$, $\lambda_1 \geq \lambda_2 \geq \ldots \geq \lambda_n$. We want to construct an expression pattern $x \in \mathbb{R}^n \backslash \{0\}$,

$$x = \sum_{i=1}^{n} c_i \, w_i \, , c_i \in \mathbb{R} \tag{21}$$

such that its Moran's I assumes some given value $M \in [-1,1]$,

$$M = M(x) = \frac{1}{|W|} \frac{x^T W x}{x^T x} = \frac{\sum_{i=1}^{n} \lambda_i \, c_i^2}{\sum_{i=1}^{n} c_i^2} \tag{22}$$

Multiplying by the denominator and sorting terms, we obtain

$$\sum_{i=1}^{n} (M - \lambda_i) \, c_i^2 = 0 \tag{23}$$

As $c_i \neq 0$ for at least one $i$, it follows that this Equation only has a solution if $M \in [\lambda_n, \lambda_1]$, and for ease of presentation, we require that $M \in (\lambda_n, \lambda_1)$. For typical neighborhood grids, $\lambda_n$ will be close to $-1$ and $\lambda_1$ will be close to $1$, so this is no serious restriction. A straightforward way to construct suitable coefficients $c_i$ is to sample $d_i \sim_{iid} \mathcal{N}(0,1)$, $i = 1, \ldots, n$. Then, let

$$\begin{aligned} s_+ &= \sum_{i, M - \lambda_i > 0} (M - \lambda_i) d_i^2 \\ s_- &= \sum_{i, M - \lambda_i \leq 0} (M - \lambda_i) d_i^2 \end{aligned} \tag{24}$$

and verify by elementary calculations that the coefficients

$$c_i = \begin{cases} d_i / \sqrt{s_+} & \text{if } M - \lambda_i > 0 \\ d_i / \sqrt{s_-} & \text{if } M - \lambda_i \leq 0 \end{cases} \quad , i = 1, \ldots, n \tag{25}$$

satisfies Equation 23. We finally shift $x$ to mean zero and scale it to unit variance for better visualisation.

In the simulation for Main Figure 1B, we chose a quadratic $50 \times 50$ grid with each point having exactly four neighbours (top, bottom, left, right). Additionally, the leftmost spots shown in the pattern connect to the rightmost spots, and the top spots are neighbours to the bottom spots. In other words, the spots form a torus. On this grid, Moran's I is invariant under right and up shifts of patterns and therefore allows us to generate more diverse patterns by translocating them by a random horizontal and vertical offset.

### Anticorrelation of non-spatial genes with coverage pattern

We observed that the coverage pattern (i.e., the library size per spot, the total read counts mapped to each spot) typically shows a strong spatial structure of the coverage pattern (Supplementary Figure S1C). As different cell types are likely to contain different amounts of RNA, this is to be expected for most tissues, leading to cell-type specific, spatial variation of library size per spot. Consequently, SVG expression patterns often correlate or anticorrelate with this coverage pattern. This can lead to severe artefacts when simulating count numbers of random genes by shuffling/resampling count numbers from an observed gene pattern:

Let us consider a gene which follows the total spot coverage pattern and let us randomly permute its count values. If $N = (N_s)$ is the total coverage count pattern ($s$ ranges across all spots), and $c = (c_s)$ is the count pattern of this gene, let $\tilde{c} = \left(c_{\pi(s)}\right)$ the permuted pattern, where $\pi$ is a random permutation of the spots. The relative abundances $\lambda = (\lambda_s)$ respectively $\tilde{\lambda} = \left(\tilde{\lambda}_s\right)$ can be calculated from this as

$$\tilde{\lambda}_s = \frac{c_{\pi(s)}}{N_s} = \lambda_{\pi(s)} N_{\pi(s)} \cdot \frac{1}{N_s} \tag{26}$$

Note that $\left(\lambda_{\pi(s)} N_{\pi(s)}\right)$ is a vector with positive entries which is uncorrelated to $\lambda_s$ (due to the permutation $\pi$). This means that if $\lambda_s$ is correlated to $N_s$, it will be anticorrelated to $\frac{1}{N_s}$. Consequently, $\tilde{\lambda}_s$ will be anticorrelated to $\lambda_s$. This artefact is dangerous because patterns that are considered to have no spatial structure unintentionally inherit spatial structure from the total coverage pattern $N$. We note that this artefact could be the reason for the excessive number of "significant" SVGs found by some methods that do not adequately correct for the spatial structure of the count pattern as a confounding factor.

### Coverage-adjusted local resampling

For sensitivity and specificity assessment of our method, as well as for checking its robustness against model violations, we need to generate realistic spatial sequencing data with tuneable degrees of non-spatial noise applied to observed expression patterns. To do so, we developed a coverage-adjusted local resampling method. The basic idea is to take an initial pattern and resample, for each spot, its expression value from the set of its k nearest neighbours. Here, $k$ can be chosen from $k = 0$ (no changes), $k = 1$ (sampling from nearest neighbours) up to $k = \infty$ (random resampling from all available spots). According to the previous section, a naive implementation would not lead to the goal; on the contrary, we would generate spatially structured patterns that would anticorrelate with the total count pattern. We have therefore developed a method that works with relative frequencies instead of counts, while preserving the total coverage pattern as much as possible.

With the notation from above, let $\left(\lambda_s^g\right)$ the matrix of relative abundances, $\lambda_s^g = \frac{c_s^g}{N_s}$, with $c_s^g$ the count number of feature $g$ in spot $s$. Pick a gene/feature $x$ with relative abundance pattern $\lambda^x = (\lambda_s^x)$. Let $\lambda^y = \left(\lambda_s^y\right)_{s=1,\dots,n}$ a "local" bootstrap sample of $\lambda^x = (\lambda_s^x)$. This means that for each spot $s$, the value $\lambda_s^y$ is sampled uniformly at random from $\{\lambda_t^x; t \in N(s)\}$, where $N(s)$ is a specified neighbourhood of $s$, e.g., a circle of fixed radius around $s$ (Supplemental Figure S1C and D). Specifically, for $N(s) = \{1, \dots, n\}$, we perform an unrestricted bootstrap which corresponds to erasing all spatial information potentially contained in $\lambda^x$. We want to add a perturbed twin $y$ of $x$ to the dataset. This is achieved by letting the relative abundances be

$$\mu_s^g = \frac{\lambda_s^g}{1 + \lambda_s^y} \quad , \; g \in \{1, \dots, p\} \cup \{y\} \tag{27}$$

and the library sizes per spot

$$M_s = \left(1 + \lambda_s^y\right) \cdot N_s \tag{28}$$

This leads to the updated count matrix

$$b_s^g = \mu_s^g M_s = \begin{cases} \dfrac{\lambda_s^y}{1 + \lambda_s^y} N_s & g = y \\ c_s^g & g \neq y \end{cases} \tag{29}$$

Note that $b_s^y$ does not need to be an integer. As some methods require integer counts as input, and to avoid bias introduced by rounding of low abundance expression levels, we replace $b_s^y$ with $\lfloor b_s^y \rfloor + \epsilon_s^y$, where $\lfloor b_s^y \rfloor$ is the largest integer not greater than $b_s^y$, and $\epsilon_s^y \sim \text{Bernoulli}\left(b_s^y \bmod 1\right)$ is a Bernoulli random variable that takes the value 1 with probability equal to the modulus of $b_s^y$.

### p-value calibration

We checked the calibration of SpaCo's SVG test (i.e. uniform distribution of p-values under the null hypothesis) and another state-of-the-art method, SPARKX[4]. Note that the coefficients $c_i$ in SPACO's test statistic (Equation 16) depend on the entire data set. Therefore, we validated the SVG test in the anterior brain dataset and in the liver dataset separately. We calculated the respective coefficients $c_i$ and sampled $10^5$ expression patterns from our null model (independent, standard normally distributed values per spot). After calculating the corresponding p-values, we compared the empirical p-value distribution with the expected (uniform) p-value distribution, both for SpaCo and for SPARKX[4].

To further check robustness against violations of the null model, we applied the same procedure to $10^5$ random patterns obtained from the spatial genes Gpr88 (brain) and Cyp2e1 (liver) by coverage-adjusted, unrestricted resampling. Main Figure 1C confirms that both methods produce well-calibrated p-values in all four scenarios.

Each split dataset underwent entirely independent processing, and the adjusted Rand index and normalised mutual information were used to assess the consistency of the two resulting clusterings. SPACO displayed better reproducibility in the brain data (Figure 1C).

### Split dataset analysis and clustering comparison

For assessing the consistency of clustering, each of the three datasets was divided into two by splitting the spot grid into pairs of mutually adjacent spots that are shifted horizontally by one unit (Supplementary Figure S1A). The left (right) spot of each pair was assigned to grid 1 (grid 2). Some points at the left/right margins that did not have a partner were discarded. This resulted in a pair of artificial datasets for each sample. The liver datasets had 777 spots per group, the anterior mouse brain dataset 1319 spots per group, and the posterior mouse brain dataset 1652 spots per group. Each of these datasets was then processed independently using either the Seurat (V. 4.3.0.1) standard pipeline for 10X Visium spatial data analysis (Satija, 2023) or the standard SPACO analysis pipeline. Upon completion of processing, the calculated significant spatial components were integrated into the pre-existing Seurat objects via Seurat's *CreateDimReducObject* function. For contrasting the spatial component clustering result with the PCA-based clustering, PCA was performed on the two datasets per sample utilising the *RunPCA* function of Seurat. The number of principal components employed for clustering was determined through an elbow plot of the principal components. The neighbourhood graph for clustering was computed with the *FindNeighbors* function, using either the significant principal components or spatial components. Louvain clustering was executed through the Seurat's *FindClusters* function. To ensure a fair comparison, the resolution parameters were modified to yield the same number of clusters for both original datasets and both methods (the agreement of clusterings becomes increasingly fragile with increasing number of predicted clusters). The clustering agreement between two split datasets was measured by the adjusted Rand index (ARI) and the normalized mutual information (NMI). ARI was calculated using the mclust (V. 6.0.0) package's *adjustedRandIndex* function (Scrucca et al., 2016). NMI was calculated with the aricode (V. 1.0.2.) package, using the "max" variant of the *NMI* function (CRAN - Package aricode, n.d.). As Louvain clustering starts with a random initiation, the process was repeated with 1000 different random seeds. Results illustrated in Figure 1C are derived only from those paired datasets that resulted in the identical number of clusters for both splits.

### Assessment of the denoising property of the SpaCo projection

For the evaluation shown in Figure 2A we performed coverage adjusted local resampling of the datasets for the depicted radii. All patterns were projected $L$-orthogonally onto the relevant SpaCs. The delta values in the right middle figure were calculated as the squared Euclidean distance between the original pattern and the denoised pattern, divided by the number of spots (regarding the denoised pattern as an approximation of the original, this is the average squared residual per spot). For the figure on the bottom right, we calculated the Euclidean distance of all SVG per radius from the resampled gene to the original and from the denoised gene to the original.

### Spatially variable gene detection benchmarking

To assess the capacity of SPARKX and SPACO to identify spatially variable genes at various noise levels, we used a gold standard of 757 spatial marker genes that have been previously documented for the mouse brain anterior dataset (Bae et al., 2022), available in the supplementary data. For each gene in this collection, we added a non-spatial 'twin' gene as a negative control, which was obtained from the gene's expression pattern by coverage-adjusted, unrestricted resampling. All other genes merely served as background data. Subsequently we increased the difficulty of the SVG detection task step by step and generated ten individual benchmarking datasets that were identical to the previous one, except for the 757 marker genes. The spatial pattern of those was blurred by coverage-adjusted local resampling, for increasing radii (r = 1, 5, 20, 40, see Figure 1E). Using the 757 marker genes as cases and their randomly permuted twins as controls, we calculated a Receiver Operating Characteristic (ROC) curves in each dataset based on the SVG test of SPACO and SPARKX. For each scenario and each test method, we generated 10 independent datasets. We report a median ROC curve for each scenario and method using the median of the 10 sensitivity values for a given specificity. For SVG testing with SPARKX, we followed the protocol in (Zhu et al., 2021). In particular, all genes with zero counts were removed, the coordinates were normalised to a mean of 0 and a variance of 1, and the genes were similarly adjusted as described in the methods section of SPARKX. The 'mixture' option was selected for kernel choice.

### Experimental data analysis

10X Visium brain datasets

The Visium spatial transcriptomics dataset for the mouse brain was downloaded from the 10X Genomics Data Repository. The mouse brain serial anterior section slide contains 2695 spots and 32,285 genes, and the posterior slide contains 3353 spots and 31,053 genes. Sequencing metrics can be found in the 10X Genomics Data repository https://www.10xgenomics.com/resources/datasets [8].

10X Genomics Visium liver data

The spatial transcriptomics dataset for young mouse liver was taken from Nikopoulou et al., 2021. It contains 1596 spots and 32285 genes. Sequencing metrics and sample preparation details can be found in the original publication.

The GO-Term analysis shown in Figures 2 B and C and supplementary figure S1E was done using Clusterprofiler (V. 4.2.2.) [10] using the standard parameters and "biological process" as Term input. Gene lists for the comparison of spatially variable genes detected using SPACO and described in the literature were obtained from Bae et al., 2022 for the brain dataset and from Hildebrandt et al., 2021 for the liver dataset.

### Supplemental Figures

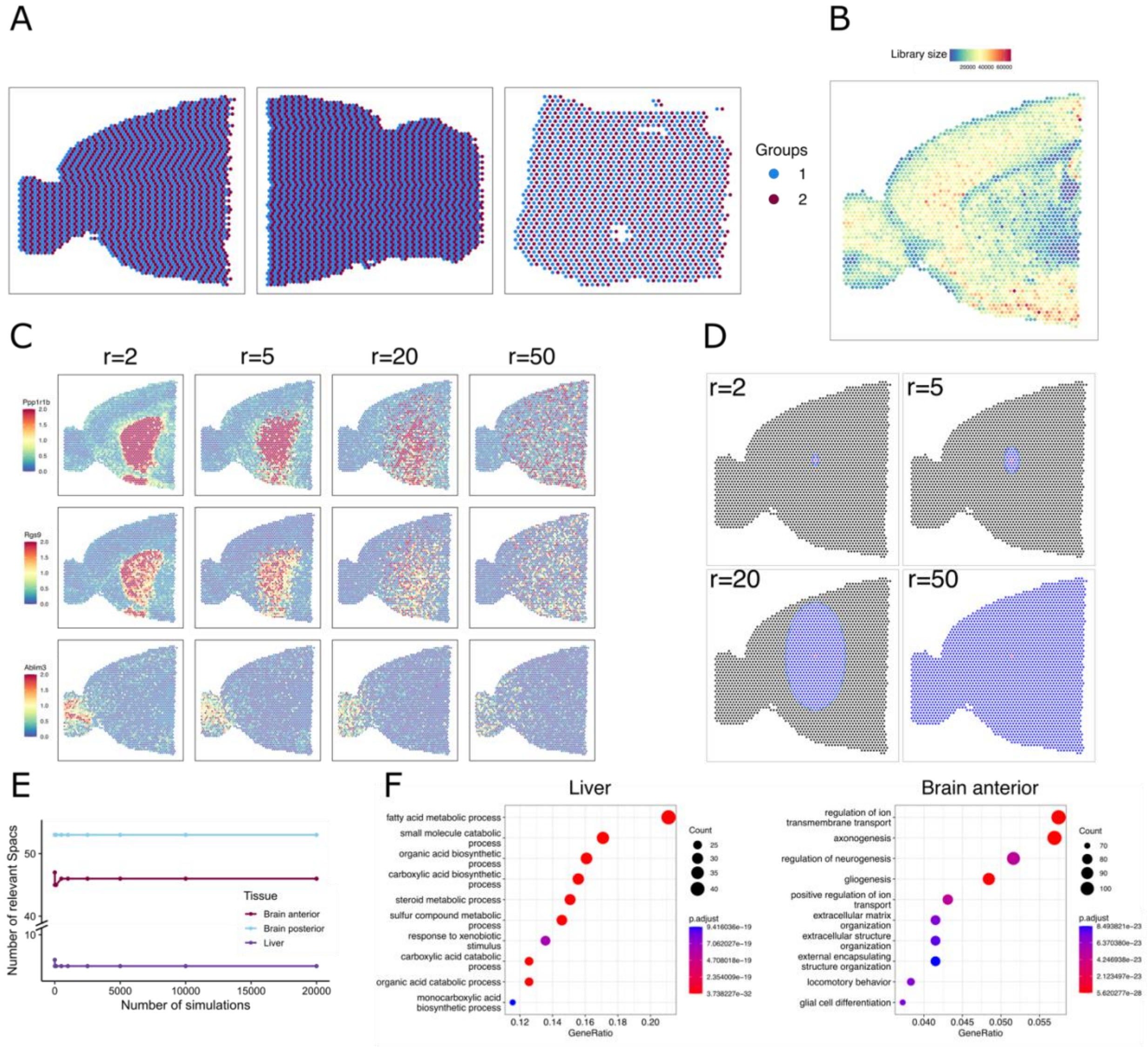


**Supplemental Figure S1.** *A) Spot grid of the murine anterior & posterior brain (left, middle) and the liver dataset (right) used for clustering consistency evaluation. Two artificial datasets were generated by splitting the grid into two groups (blue and red spots). Each blue spot is paired to the red spot shifted by one unit to the right. Spot pairs are considered representing identical cell types for comparison purposes. B) Raw coverage feature plot of the mouse anterior brain section used in the manuscript for method evaluation. C) Normalised gene expression of three representative genes (Ppp1r1b, Rgs9 and Ablim3) over increasing noise levels. The R-value indicates the neighbourhood radius from which the expression values were resampled. D) Visualisation of the neighbourhood size corresponding to their radius. The spot in red is the spot to be resampled; blue cells represent the corresponding sampling pool. E) Number of relevant spatial components k picked as the k-the eigenvector below the 5% quantile of the smallest eigenvalue occurring in a data set,* $X'$ *which the spots of* $X$ *(the original data) were permuted randomly. On the x-axis number of simulations used for relevant SpaC computation. E) GO-Terms of SVG determined with SPACO not listed as spatially regulated for the murine liver and the anterior brain dataset.*
